# Distinct polysaccharide growth profiles of human intestinal *Prevotella copri* isolates

**DOI:** 10.1101/750802

**Authors:** Hannah Fehlner-Peach, Cara Magnabosco, Varsha Raghavan, Jose U. Scher, Adrian Tett, Laura M. Cox, Claire Gottsegen, Aaron Watters, John D. Wiltshire-Gordon, Nicola Segata, Richard Bonneau, Dan R. Littman

## Abstract

The gut-dwelling *Prevotella copri*, the most prevalent *Prevotella* species in human gut, have been associated with diet as well as disease, but our understanding of their diversity and functional roles remains rudimentary, as studies have been limited to 16S and metagenomic surveys and experiments using a single type strain. Here, we describe the genomic diversity of 83 *P. copri* isolates from 11 donors. We demonstrate that genomically distinct isolates, which can be categorized into different *P. copri* complex clades, utilize defined sets of plant polysaccharides that can be predicted in part from their genomes and from metagenomic data. This study thus reveals both genomic and functional differences in polysaccharide utilization between several novel human intestinal *P. copri* strains.

## Introduction

Humans live in close association with a teeming bacterial ecosystem known as the gut microbiome. The diverse inhabitants of this extraordinary environment perform beneficial and remarkable services for their host, out-competing harmful bacteria and digesting the majority of dietary fiber (El Kaoutari et al., 2013; Martens et al., 2008). The relative abundance of two genera, *Bacteroides* and *Prevotella*, is inversely correlated in the intestine (Arumugam et al., 2011; Costea et al., 2018). Although a *Bacteroides*-dominated microbiome is more common, *Prevotella* are present at >10% relative abundance in the stool of 10-25% of healthy American and European individuals (Koren et al., 2013). Compared to at least 25 species of *Prevotella* in the oral cavity, and at least 17 species of intestinal *Bacteroides*, only five *Prevotella* species have been reported in human intestine, with *Prevotella copri* being the most abundant (Accetto and Avgustin, 2015; Ferrocino et al., 2015; Ibrahim et al., 2017; Li et al., 2009; Lin et al., 2013; Liu et al., 2018). Although mechanistic models of metabolism, growth, and colonization have been established for several intestinal *Bacteroides* and oral *Prevotella*, studies of intestinal *Prevotella* have lacked experimental tools and relied heavily on only two strains (Hayashi et al., 2007).

Many researchers have observed *Prevotella*’s presence and abundance by 16S sequencing, but its genomic diversity and biological properties have not been described. Because *Prevotella copri* are highly sensitive to oxygen, and difficult to isolate and cultivate, previous studies have been limited to the single commercially available type strain, DSM 18205, to perform metagenomic comparisons and draw experimental conclusions. *Prevotella* spp. have extensive gene acquisition and loss (Gupta et al., 2015; Zhu et al., 2015). In fact, accessory genes within the *P. copri* DSM 18205 genome account for around 40% of all genes encoded within the genome, the highest proportion of accessory genes identified in a comparison of 11 gut-associated species (Zhu et al., 2015). This estimate should be revised, however, since a metagenomic study (supplemented by whole genome sequencing data from the present study) affirms that *P. copri* is diverse, comprising four genomically distinct clades (Tett et al., 2019). Further improvements in genomic comparisons of *P. copri* may be expected soon, since another study reports a closed genome of *P. copri*, generated from long-read sequencing of whole microbiomes (Moss and Bhatt, 2018). Whether the genomic diversity in human *P. copri* translates to functional diversity has not yet been fully explored. This is particularly relevant in light of elevated frequency of *P. copri* colonization in some cohorts of treatment-naïve rheumatoid arthritis patients (Maeda et al., 2016; Scher et al., 2013). A better understanding of whether there are functional differences in strains from patients compared to healthy subjects will contribute to determining whether there is a causal relationship between *P. copri* and this autoimmune disease.

Several studies have found associations between diets and the presence of intestinal *Prevotella*. *Prevotella* have been positively associated with diets high in plant fiber and carbohydrates, and negatively associated with fat and amino acid diets (De Filippo et al., 2010; Ruengsomwong S, 2016; Wu et al., 2011). Genes associated with carbohydrate catabolism were identified in *P. copri* metagenomes and correlated with vegan diets (De Filippis et al., 2019). The plant polysaccharide xylan has been used to select for growth of human *Prevotella* (Tan et al., 2018). *Prevotella* isolates from livestock grow in the presence of plant polysaccharides and encode gene products capable of breaking down these polysaccharides (Accetto and Avgustin, 2019). Thus, diet may be an important factor promoting growth of intestinal *Prevotella*. Whether human *P. copri* are uniquely suited to digest dietary components highly abundant in plant-based diets remains unsubstantiated by experimental and genomic evidence.

Intestinal commensal bacterial genomes encode a system of genes, often clustered, that collaborate to harness energy from carbohydrates present in the host’s intestinal lumen. Extensive and careful work in *Bacteroides* species has identified several polysaccharide utilization loci (PULs) that are specific for glycans derived from plant, animal, and host sources (Martens et al., 2008; Martens et al., 2011). Bacteria lacking one of several critical genes in any polysaccharide-specific PUL fail to grow when the polysaccharide is provided as the sole carbon source (Martens et al., 2011; Raghavan et al., 2014). PULs have been linked to diet and subsequently exploited in experimental systems. Notably, a seaweed-utilizing PUL was horizontally transferred from marine bacteria to intestinal bacteria of seaweed-ingesting hosts (Hehemann et al., 2010). This PUL was experimentally introduced to *B. thetaiotaomicron*, and seaweed-selective growth was demonstrated in laboratory mice (Shepherd et al., 2018). Although by far the most extensive work on PULs has been performed in *Bacteroides* species, PULs have been annotated in *Prevotella* type strains and used to predict growth on plant polysaccharides in *Prevotella* isolates from livestock (Accetto and Avgustin, 2015, 2019). Despite their association with plant diet and PUL predictions, the functional capabilities of human intestinal *Prevotella* have not been tested, due to a dearth of cultured human intestinal *Prevotella* isolates.

Previous studies have been limited to analysis and experimentation with the available genomes and strains of *Prevotella* species found in human, two of which were isolated from the same individual (Hayashi et al., 2007). Here, we report whole genome sequencing of 83 *Prevotella copri* isolates from the stool of eleven individuals. We observe extensive genomic diversity of *P. copri* isolates within and between hosts. This genomic variation was exemplified by specific gene clusters and linked to functional growth of isolates on predicted plant polysaccharides, explaining the association of colonization of *Prevotella* with a plant-rich diet.

## Results

### Isolation of *Prevotella* from human stool

To investigate the genomic diversity of *Prevotella copri* and also identify strains for functional analysis, we developed a qPCR screening and isolation strategy for *P. copri* from human stool (Figure S1, Table S1, **Methods**). *P. copri* was identified in 31/63 individuals by qPCR (Figure S1). 16S rRNA gene sequencing of the hypervariable V3-V4 region found *P. copri* in 41/63 individuals, with 18/63 individuals harboring *P. copri* at levels >10% relative abundance (Figure 1A). Relative abundance ranged from < 0.01 to 0.766. From the V3-V4 16S rRNA gene data, five *P. copri* 16S variants were identified (Figure 1A, **Supplemental file 1**). The *P. copri* 16S variants displayed 2-6 single-nucleotide variants (SNVs) within the 253 bp region, corresponding to >97% sequence-sequence identity (Callahan et al., 2016). *P. copri* variants 1 and 2 were the most abundant ones in the stool samples. *P. copri* detection by qPCR positively correlated with *P. copri* detection by 16S sequencing (Figure 1B). The qPCR screen reliably detected *P. copri* in samples with >1% *P. copri*, as determined by 16S sequencing, but was less sensitive for samples containing *P. copri* at lower relative abundance. We collected 83 isolates from eleven individuals harboring *P. copri* (Table 1), one of whom had <1% *P. copri* (**Methods**).

**Figure 1.**
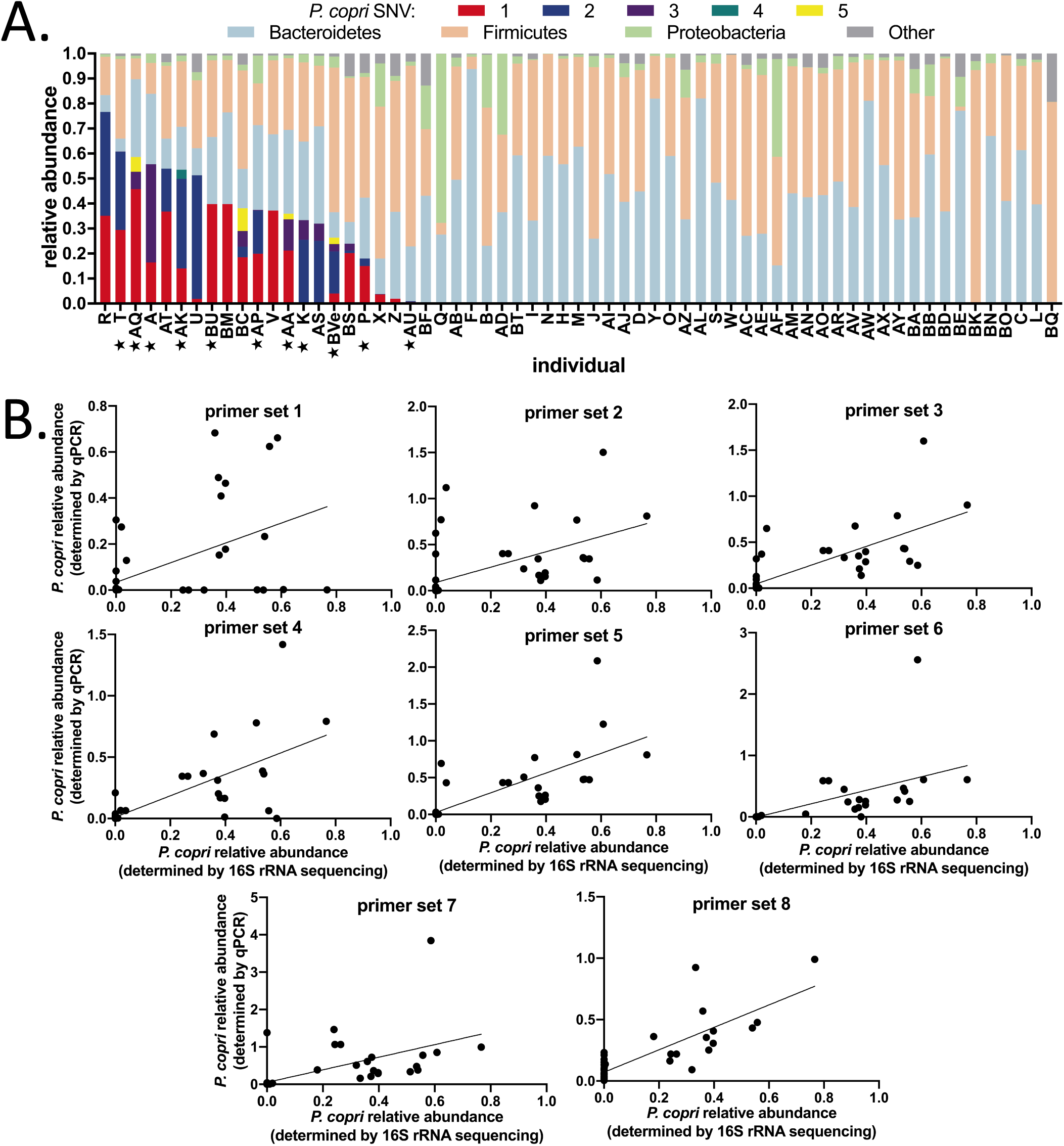
Identification of *Prevotella copri* in human stool. **A.** Relative abundance of *Prevotella copri* 16S SNVs in human stool quantified by 16S sequencing. Percentage of total reads from 16S rRNA gene sequencing is shown. Stars indicate individuals from whom *P. copri* was isolated. **B.** Correlation of *P. copri* abundance determined by 16S sequencing and qPCR using each qPCR primer set (Table S1).

**Table 1.**
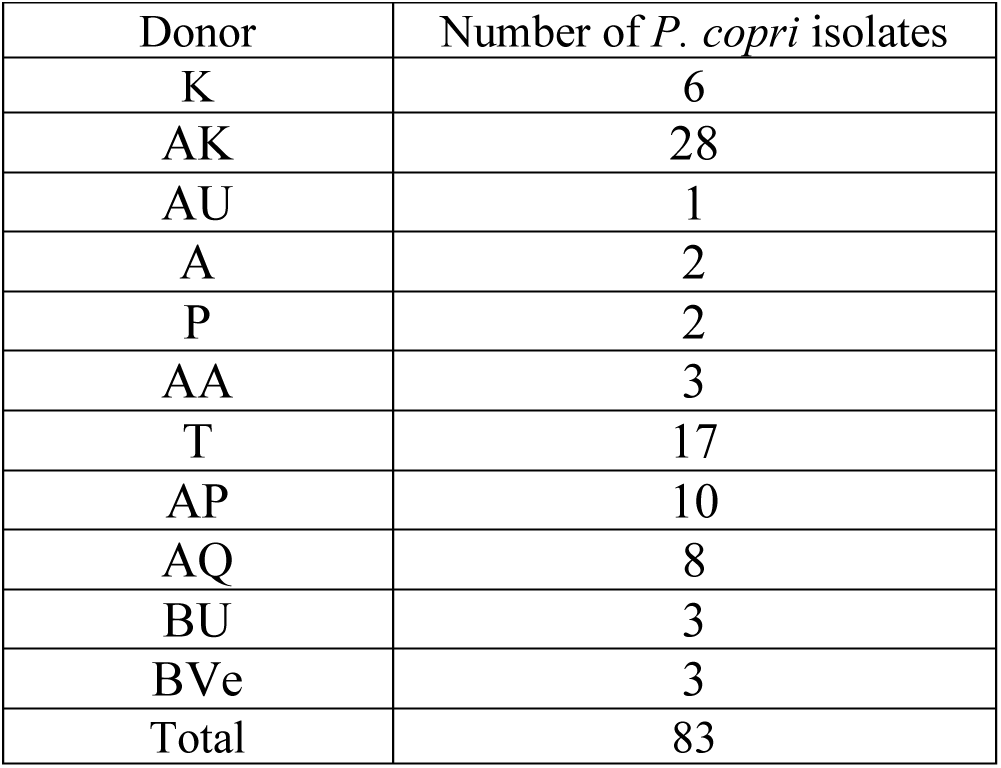
Number of Prevotella copri isolates collected from individual donors.

### Variability in *P. copri* isolates

Our earlier study indicated differences in the *P. copri* genomes from new onset rheumatoid arthritis and healthy individuals (Scher et al., 2013). Other studies have since described genetic variation between strains from healthy individuals (De Filippis et al., 2019; Zhao et al., 2019). We assessed whether genetic diversity existed between *P. copri* isolates from different individuals, or, furthermore, between isolates from the same individual, by sequencing the full genomes of the isolates. Alignment of filtered isolate sequencing reads to the *P. copri* PanPhlAn database revealed that fewer than 60% of reads mapped to the reference genome, indicating unexpected and unexplored diversity of *P. copri* genomes in our isolates (Figure 2A) (Scholz et al., 2016).

**Figure 2.**
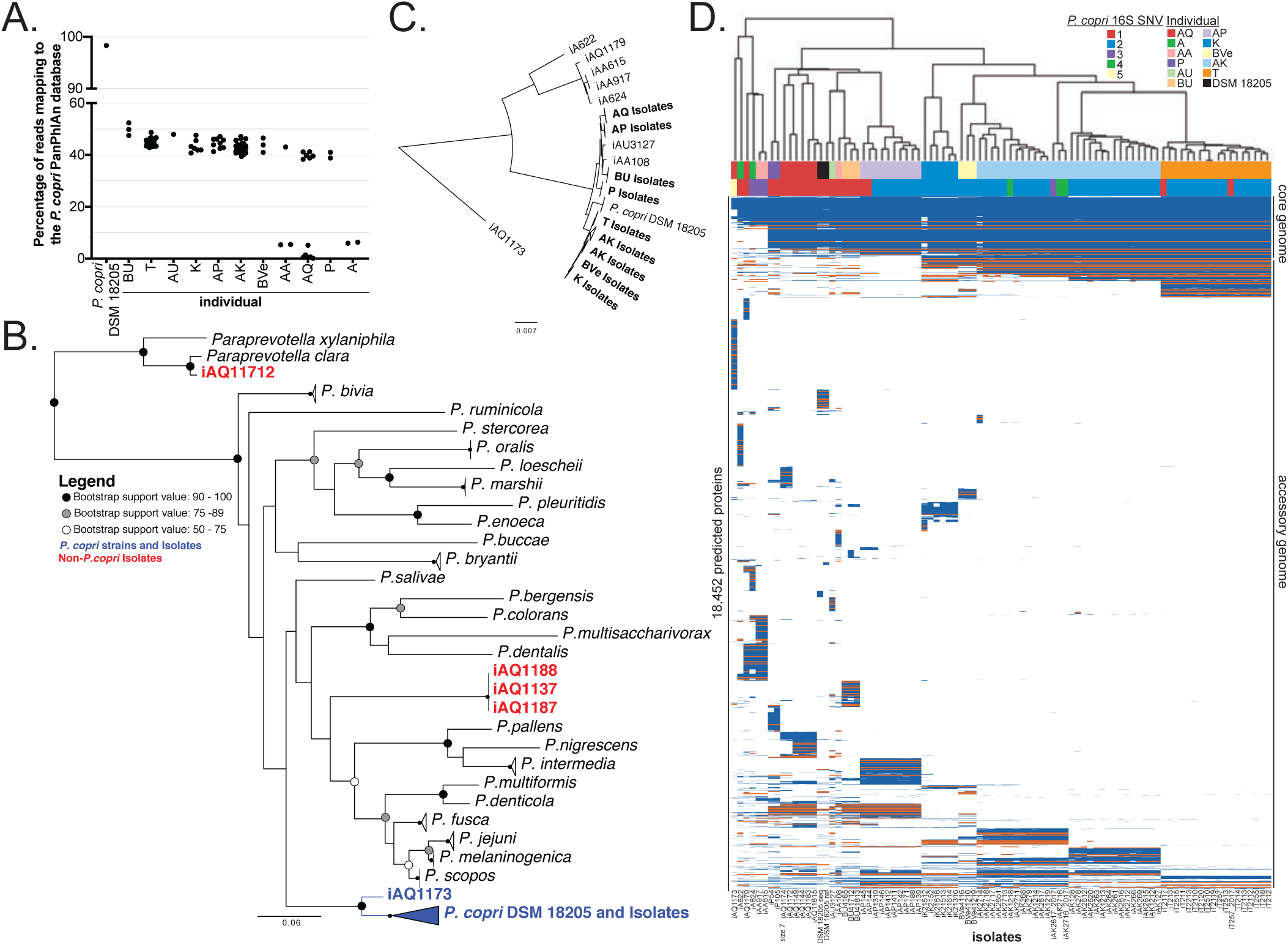
Hidden diversity in *Prevotella copri* whole genome sequences. **A.** Alignment of whole genome sequencing reads of each isolate to the *P. copri* PanPhlAn database. Each point represents alignment of a single isolate genome. **B.** Phylogenetic tree based on the 16S rRNA gene sequences. *P. copri* isolates are shown in blue. Three *Prevotella* isolates and one *Paraprevotella* isolate that were not *P. copri* are shown in red. These isolates were excluded from further analyses. **C.** Phylogenetic tree based on the concatenated alignment of 30 conserved ribosomal protein gene sequences. **D.** Heatmap of *P. copri* isolate whole genomes showing presence and absence of predicted proteins, and highlighting *susC* genes. Predicted proteins were clustered at 90% amino acid identity. Blue = present; white = absent; orange = *susC*. Isolates from the same donor are indicated by color in line 1 above the heatmap, as well as by the first letters of the isolate name below the heatmap.

Evaluation of the *de novo* assembled genomes revealed low contamination and similar genome size among all isolates (Table S2) (Bankevich et al., 2012; Parks et al., 2015). Phylogenetic analysis based on the 16S rRNA gene revealed that most of the isolates clustered with *Prevotella copri* DSM 18205 (Figure 2B). One isolate clustered with *Paraprevotella*, and three isolates clustered with other species of *Prevotella*; these four isolates were excluded from the rest of the analysis. A separate phylogenetic analysis based on 30 conserved ribosomal proteins further confirmed that the isolates clustered with *P. copri*, and also revealed distinct clades (Figure 2C). Comparison with a catalog of *P. copri* genomes from metagenomic assembly confirmed that most isolates were in a main clade (clade A), four isolates were in another clade (clade C), and one isolate each was in two additional clades (clades B and D) (Tett et al., 2019). Hosts AQ, A, and AA harbored isolates belonging to more than one clade (Figure 2C).

### *P. copri* genomic diversity between hosts

Despite close phylogenetic placement, most isolates exhibited striking genetic dissimilarities to the *P. copri* reference strain (Figure 2A). To better understand the differences in the *P. copri* genomes, we compared the presence and absence of predicted ORFs in all the isolates (Figure 2D). This analysis revealed a core genome of approximately 1750 genes, suggesting accessory genomes ranging from approximately 1250-2250 genes per isolate.

Clustering of the isolates by presence/absence of predicted ORFs revealed that isolates often clustered by donor. When isolates from the same donor fell into the same clade, accessory genomes were often unique to isolates from individual hosts, as visualized by blocks of genes in the accessory genomes in Figure 2D. Host AK had isolates with distinct accessory genomes within the same clade (Figures 2C and 2D). Annotation of the genomes revealed that, among other functions, many of the host-specific genes were involved in polysaccharide transport, as indicated by the high frequency of *susC* genes (Figure 2D).

### Polysaccharide utilization prediction

SusC is the bacterial transmembrane protein that transports polysaccharides into the periplasm. SusC collaborates with the outer-membrane glycan-binding protein SusD, as well as other proteins, to bind and digest polysaccharides. Conveniently, the genes for this polysaccharide utilization machinery usually occur grouped in close proximity to one another in the genome in polysaccharide utilization loci (PULs) in Bacteroidetes, although occasionally genes encoding polysaccharide catabolism enzymes are located elsewhere in the genome (Sonnenburg et al., 2010). We hypothesized that the genetic diversity of *susC* genes might indicate functional diversity in polysaccharide utilization by the isolates. We surveyed the *susC* and surrounding genes in the isolate genomes to predict the repertoire of PULs and, by extension, polysaccharide substrates for the *P. copri* isolates.

By clustering *susC* genes at 90% identity, and examining the neighboring gene products within each genome, we identified 125 *susC* gene clusters with a neighboring *susD* gene, which we called predicted PULs (pPULs) (**Methods**, Figure S2, **Table S3**). Based on the genes flanking *susC* in each pPUL, we predicted the substrate for 87 pPULs. Due to a high frequency of hypothetical proteins, we were unable to predict substrates for 38 pPULs, which were classified as “PULs of unknown substrate.” The predicted substrate, as well as the distribution within isolate genomes, was analyzed for each pPUL (Figure 3A). The same substrate was often predicted to be targeted by PULs with different *susC* clusters in different isolate genomes. For example, xylan was the pPUL substrate for *susC* clusters 2, 65, and 3. *SusC* cluster 2 was only in isolate genomes from individual BU, *susC* cluster 65 was only in the genome of isolate A622, and *susC* cluster 3 was only in isolate genomes from individual P. We identified unique pPULs for beta-glucanases and alpha-L-arabinofuranosidases (Figure 3B). Although beta-glucanases break down linkages in beta-glucan, and alpha-L-arabinofuranosidases break down linkages in rhamnogalacturonan and arabinan, a polysaccharide substrate containing both linkages has not been described. In contrast with several *Bacteroides* species, which catabolize polysaccharides from both animal and plant sources, all identified pPUL substrates for *P. copri* isolates were from plant sources, and none were from animal sources (Martens et al., 2011; Salyers AA, 1982). Consistent with this prediction, the V1-V3 16S sequences of select isolates were most closely related to two previously reported non-omnivore-associated *P. copri* 16S sequences (De Filippis et al., 2016) (Figure S3, **Supplemental file 2**).

**Figure 3.**
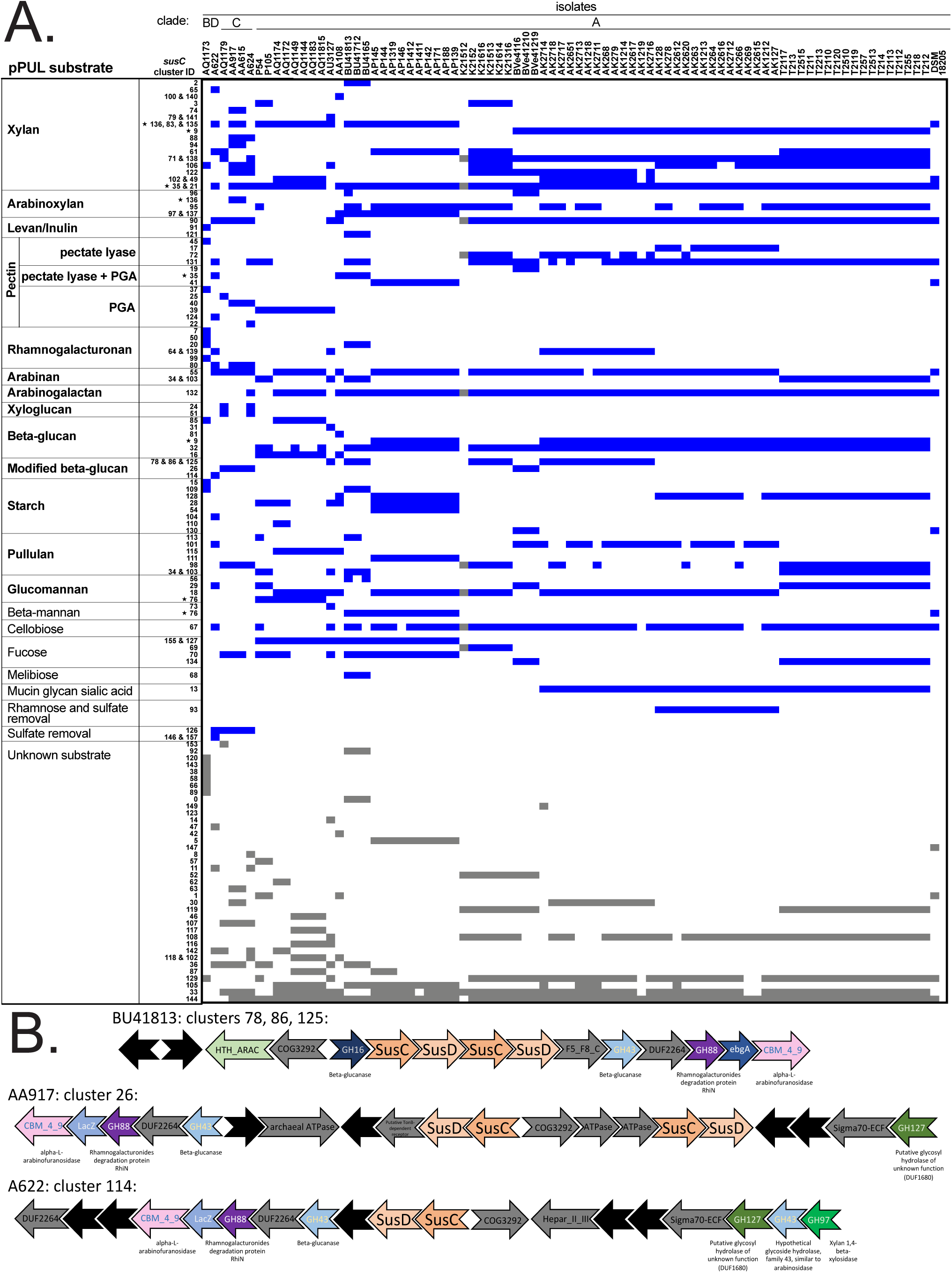
Annotation of polysaccharide utilization loci in *P. copri* isolates. **A.** PUL substrates were predicted for each *susC* cluster gene group (Figure S2, **Table S3**, **and Methods**). Predicted substrate usage for each isolate or group of isolates is indicated by a blue box. Stars next to cluster IDs indicate *susC* genes that have different surrounding enzyme genes in different isolates. **B**. Schematic of modified beta-glucan pPULs (polysaccharide source not yet identified). Protein families are labeled, and specific predicted functions are listed beneath relevant genes. Hypothetical proteins are unlabeled in black. Genes unlikely to be relevant to polysaccharide utilization are labeled and colored gray.

Consistent with the 17 predicted PULs for *P. copri* DSM 18205, the number of predicted PULs per isolate ranged from 14-28, with a median of 23 (Terrapon et al., 2018). Intestinal *Prevotella* have been reported to utilize xylan and, as expected, xylan pPULs were ubiquitously represented throughout the isolate genomes (Tan et al., 2018). A survey of the xylan pPULs in our dataset revealed additional features (Figure S4): All isolates contained xylan pPULs, as many as seven. Each genome contained 4-5 xylan catabolism enzymes that were present in pPULs. *SusC/D* pairs in xylan pPULs often occurred in tandem.

### Polysaccharide utilization *in vitro*

Based on the pPUL analysis, we predicted that *P. copri* isolates would grow in the presence of plant polysaccharides identified in the pPUL substrate repertoire (Figure 3A). We therefore tested the ability of *P. copri* isolates to use predicted polysaccharides as the sole carbon source *in vitro* (Table S4, substrates in bold in Figure 3A). Growth on individual polysaccharide substrates revealed functional diversity between isolates (Figure 4A, 4B, Figure S5). For example, isolate AA108 reached a high OD when pullulan was provided as the sole carbon source, but isolates P54 and T2112 did not. Interestingly, isolates from the same individual grew in the presence of different polysaccharides: compared to isolate AK1212, isolate AK2718 grew to a higher OD in the presence of PGA and rhamnogalacturonan, and a lower OD in the presence of beta-glucan and starch. As predicted, none of the isolates grew when the animal polysaccharides chondroitin or heparin were provided as the sole carbon source (Figure S5). Growth (or lack of growth) on particular polysaccharides distinguished isolates from each clade (Figure 4B, Table 2). In particular, arabinan, levan, inulin, xyloglucan, beta-glucan, and glucomannan growth distinguished isolates from each clade. All isolates except AQ1173 (which fell into clade B) grew on arabinoxylan and arabinan (Figure 4A and 4B). Isolate AQ1173 also failed to grow on xyloglucan and glucomannan. Isolate A622 (which fell into clade D) failed to grow on both levan and inulin. Tested isolates from clade C failed to grow on beta-glucan, xyloglucan, glucomannan, and rhamnogalacturonan I. Tested isolates from clade A grew well on most substrates, although there was a range of growth among clade A isolates on several polysaccharide sources.

**Figure 4.**
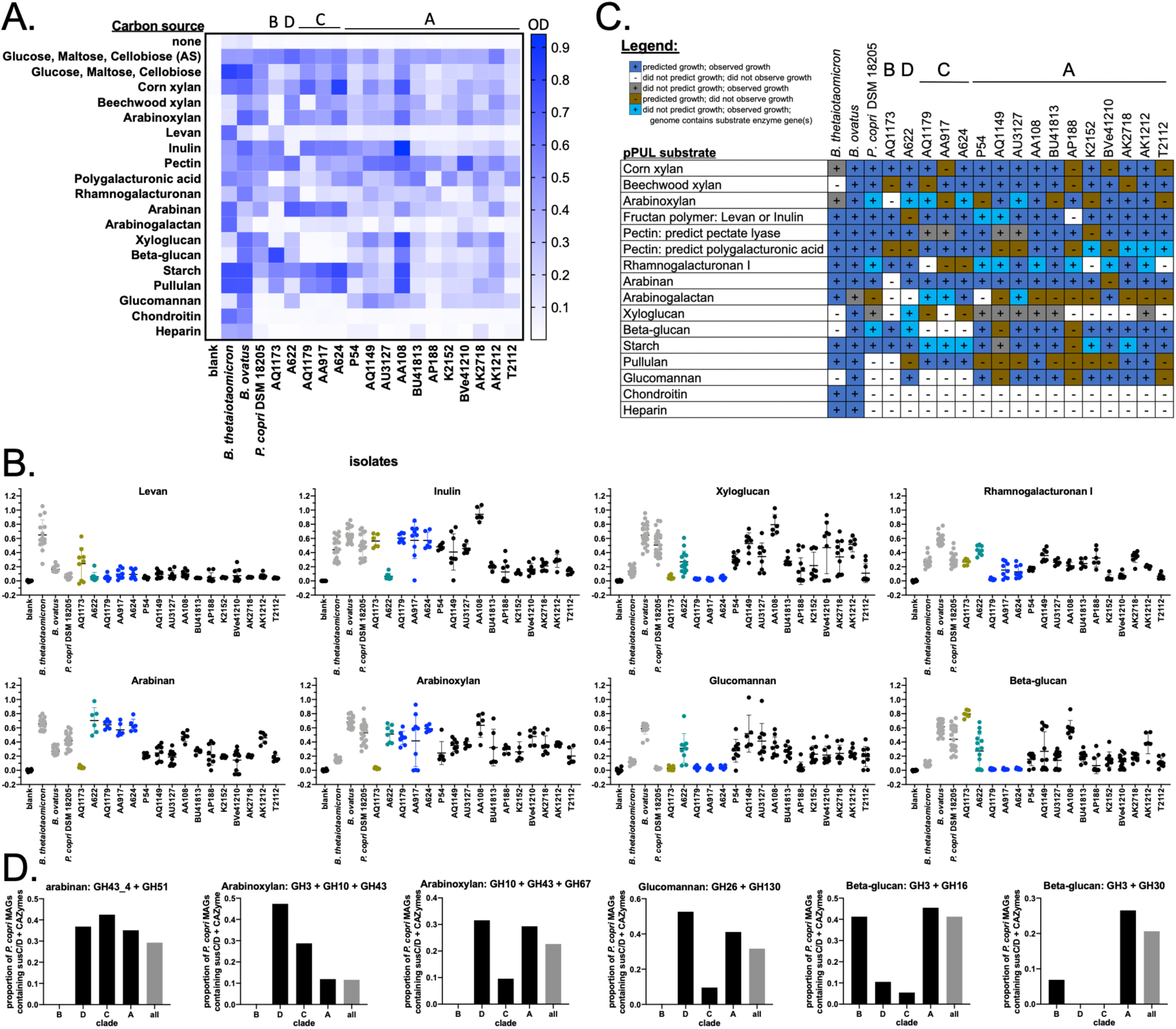
Growth of *P. copri* isolates on polysaccharides. Isolates were cultured anaerobically at 37 degrees C in YCFA media supplemented with individual polysaccharides as the sole carbon source. OD600 was measured at 24, 48, and 72h. Data represent 2-3 experiments, with each condition tested in duplicate or triplicate. The highest OD from 24, 48, or 72h for each condition for each experiment is shown. AS: media prepared by Anaerobe Systems. Heatmap (**A**) shows the mean of the highest OD for each condition of each experiment. *P. copri* complex clades are indicated above the heatmap. Graphs (**B**) show the highest measured OD on select polysaccharide substrates. Isolates from the same *P. copri* complex clade are plotted in the same color (Tett et al., 2019). **C.** Growth data were matched to growth predictions based on the presence or absence of pPULs in the genomes. Growth was considered positive if the OD was statistically significant, as determined by one-way ANOVA and post hoc Dunnett’s test using the H2O (no carbon source) as the control (**Methods**). **D.** PULs were predicted in *P. copri* MAGs. Proportion of *P. copri* MAGs within each clade containing pPULs for select polysaccharides is shown.

**Table 2.** Summary of P. copri isolate growth by P. copri complex clade.

| Substrate | clade |  |  |  |
| --- | --- | --- | --- | --- |
|  | B | D | C | A |
| Corn xylan | + | + | +/- | +/- |
| Beechwood xylan | - | + | +/- | +/- |
| Arabinoxylan | - | + | +/- | +/- |
| Levan | - | - | - | +/- |
| Inulin | + | - | + | +/- |
| Pectin | + | + | + | +/- |
| Polygalacturonic acid | - | - | + | +/- |
| Rhamnogalacturonan I | + | + | - | +/- |
| Arabinan | - | + | + | +/- |
| Arabinogalactan | - | - | + | +/- |
| Xyloglucan | - | + | - | +/- |
| Beta-glucan | + | + | - | +/- |
| Starch | + | + | + | +/- |
| Pullulan | - | - | + | +/- |
| Glucomannan | - | + | - | +/- |
| Chondroitin | - | - | - | - |
| Heparin | + | - | - | - |

Isolates AQ1173 and A622 were the only isolates that fell into clades B and D, respectively. Since growth data from single isolates limited our ability to generalize to those clades, we asked whether pPULs for relevant polysaccharides were present or absent in these clades using *P. copri* metagenome-assembled genomes (MAGs) reconstructed from metagenomic datasets (Tett et al., 2019) (**Methods**, Figure 4D). Consistent with our observation that isolate AQ1173 (Clade B) lacked arabinan, arabinoxylan, and glucomannan PULs and also did not grow in the presence of those polysaccharides (Figure 4), the Clade B *P. copri* MAGS lacked pPULs capable of degrading those polysaccharides. The Clade B MAGs did contain pPULs capable of degrading beta-glucan, however, consistent with our pPUL analysis and growth observations. pPULs were found for glucomannan and arabinan in the Clade D *P. copri* MAGs consistent with our predictions and growth observations for isolate A622 (Clade D) (Figure 4).

### Polysaccharide growth predictions

We next evaluated how well the pPULs analysis predicted *in vitro* growth: 170 isolate/polysaccharide conditions matched predictions while 85 did not (Figure 4C). We observed 42 instances in which we did not predict particular polysaccharide utilization by the isolate, yet we observed growth. For 29 of these cases of unpredicted growth, we found genes encoding relevant polysaccharide breakdown enzymes elsewhere in the genome. Perhaps the genes are part of fragmented PULs (as has been observed in *Bacteroides* fructan PULs (Sonnenburg et al., 2010)), or are simply located on a contig separate from the rest of the PUL. More puzzlingly, we observed isolates that failed to grow on a particular substrate, despite presence of a corresponding intact pPUL in the genome. For example, we predicted arabinogalactan PULs in most of genomes of clade A, yet most failed to grow on this substrate, and those that did, particularly *P. copri* isolate BVe41210, grew poorly. Perhaps *P. copri* isolates prefer a different form or source of arabinogalactan. Additionally, substrate predictions were based on gene annotations that may be incomplete or inaccurate. Overall, the pPUL analysis performed well in predicting growth. Welch’s t-test on predicted growth or unpredicted growth had a *p*-value < 0.0001 (**Methods**).

## Discussion

In this study, we observed genomic and functional diversity of human intestinal *P. copri* isolates. Whole genome sequencing exposed diversity of *P. copri* isolates, demonstrated by the low percentage of isolate whole genome reads that mapped to the *P. copri* PanPhlAn database. Within-species genomic variation, both between individuals and within individuals, was demonstrated by genetic diversity of *susC* genes and PULs and linked to functional growth of isolates on predicted plant polysaccharides. PULs predicted for several polysaccharides in large metagenomic datasets support our predictions and growth data for representative isolates, indicating that the results of this study may be generalizable beyond the isolates shown here.

### Variability in *P. copri* genomes

ORF presence/absence among isolates revealed host-specific genes in the accessory genome, several of which were *susC* transport protein genes. The variability of the *susC* genes associated with *P. copri* strains in different hosts is intriguing, given their common function of transporting glycans across the outer membrane. Since *susC* gene products are on the cell surface, they may be susceptible to selective pressure from the immune system, e.g. antibodies. Another possibility is that *susC* products from different isolates have slightly different substrates, as has been observed in two closely related *B. thetaiotaomicron* strains (Joglekar et al., 2018).

In addition to *susC* genes, we also observed integrases, transposases, and transcriptional regulators among the host-specific genes in the accessory genome. These genes may reflect the way *P. copri* behaves within the host intestine, which may be influenced by the host diet or other microbes. Isolates from host AK all fell into the same clade and possessed two distinct accessory genomes (Figure 2D), suggesting distinct niches for each type of isolate. Perhaps one type of isolate inhabits the cecum, the other the colon; alternatively, one type could reside in the lumen, while the other could be mucosa-associated. Further investigation of the genes unique to each host or isolate may reveal unique roles for individual isolates.

### *P. copri* presence in the human intestine may be diet-dependent

In humans, differences in *P. copri* 16S sequences and metagenomes have been correlated with host diet (De Filippis et al., 2019; De Filippis et al., 2016). Although our study lacks host diet information, the PUL analysis predicted and *in vitro* growth experiments confirmed that the *P. copri* isolates used only plant polysaccharides. A previous study found that particular *P. copri* V1-V3 16S sequences were associated with diet (De Filippis et al., 2016). The V1-V3 sequences of select isolates from different clades in this study were most closely related to two non-omnivore-associated sequences, consistent with their growth on plant polysaccharides (Figure S3). Although the dynamics of dietary and intestinal growth of *P. copri* within a complex microbiome require experimental validation, *in vitro* data presented here suggest that plant polysaccharides promote *P. copri* growth, but animal polysaccharides do not. These data provide a possible mechanism for the correlation between vegetable-rich diets and abundance of intestinal *Prevotella*: perhaps continuous availability of dietary plant polysaccharides is required to maintain *Prevotella* presence in the gut, and their absence may result in elimination of *Prevotella* from the human intestinal microbiome.

In hosts with more than one strain of *P. copri*, representative isolates often complemented each other to grow in the presence of more polysaccharides than any individual isolate. This point was illustrated best by isolates from donor AQ: isolates AQ1173 and AQ1179 did not grow on xyloglucan or glucomannan, but AQ1149 grew on both polysaccharides (Figure 4A). These results suggest that the total *P. copri* population in hosts with multiple strains of *P. copri* was capable of catabolizing a greater diversity of polysaccharides than any individual *P. copri* strain. Indeed, a separate study found that non-Western microbiomes contained multiple *P. copri* clades, and that some carbohydrate active enzymes were differentially represented between clades (Tett et al., 2019). Whether microbiomes of individuals with only one *P. copri* strain completely lack the ability to break down additional polysaccharides or rely on other bacterial species to maintain a full complement of polysaccharide digestion should be the subject of future metagenomic studies.

### *Prevotella* polysaccharide utilization

Although the pPUL analysis predicted growth surprisingly well, incorrect predictions may indicate interesting biology or areas in need of technical improvements. In cases in which PULs were predicted but growth was not observed, perhaps mutations in the genes encoding the enzymes or regulators have rendered the gene products non-functional. Future studies should aim to evaluate how often and how rapidly such mutations might occur in intestinal commensals. Open genomes may account for some cases in which polysaccharide utilization was not predicted by the presence of a PUL, yet growth was observed. If a PUL were split across contigs, it would be impossible to accurately predict its substrate. Recent closure of a *P. copri* genome from metagenomic analysis may help reduce such instances in future studies (Moss and Bhatt, 2018).

Utilization of xylan, which is found in cereal grains, has been repeatedly established in *Prevotella* species and in *P. copri* specifically (Accetto and Avgustin, 2019; da Silva et al., 2012; Dodd et al., 2011; Tan et al., 2018). Two distinct PULs encoding enzymes not previously associated with xylan catabolism were identified for corn xylan, birchwood xylan, and arabinoxylan in multiple *B. ovatus* and *B. xylanisolvens* strains (Rogowski et al., 2015). The numerous xylan-degrading enzymes and xylan pPULs identified in *P. copri* isolates in this study suggest that *P. copri* may have an expanded xylan degrading enzyme repertoire, and possibly a superior ability to target xylans, compared to other intestinal bacteria.

We identified three separate pPULs with enzymes capable of breaking down linkages found in beta-glucan, rhamnogalacturonan, and arabinan. The structure of the corresponding polysaccharide is mysterious. Arabinose and xylose decorate the beta-glucan backbone of xyloglucan. Accordingly, xyloglucan degrading PULs with beta-glucan, xylan, and arabinan catabolizing enzymes are well described in *Bacteroides* (Larsbrink et al., 2014; Nishinari et al., 2007). Structures and corresponding enzymes for beta-glucan, arabinan, and rhamnose, but not xylose have not yet been described. Arabinose and rhamnose cleavage enzymes imply that the PUL might act on a complex pectin, although beta-glucan linkages have not been described in pectins. One possibility is that some of the genes in these pPULs are non-functional. A more intriguing possibility is that these pPULs are unique to *Prevotella* and utilize a yet undescribed polysaccharide.

Xyloglucan is an abundant polysaccharide in vascularized plants, seeds, and food additives, yet xyloglucan utilization by known intestinal microbes is relatively rare, demonstrated only in a low proportion of *Bacteroides* species (Larsbrink et al., 2014; Nishinari et al., 2007). Despite its abundance in the human diet, xyloglucan utilization was overwhelmingly difficult to predict (Figure 4C). In fact, we observed the keystone xyloglucan endo-beta-1,4-glucanase enzyme in pPULs only in isolates A624 and AQ1179 which did not grow in the presence of xyloglucan (Larsbrink et al., 2014). The same enzyme was encoded in the genomes of isolates AA615, AA917, and A622, although not in close proximity to *susC* and *susD* genes. All other genomes lacked this enzyme. Perhaps *Prevotella* possess novel xyloglucan PUL structures or enzymes for this abundant dietary polysaccharide. Given the rarity of xyloglucan catabolism genes reported to date in *Bacteroides*, perhaps *P. copri* is responsible for xyloglucan degradation in hosts lacking *Bacteroides* species possessing xyloglucan-breakdown genes.

### Implications

The collection of *P. copri* isolates and their corresponding polysaccharide preferences described herein enable future experimentation *in vitro* and *in vivo* with isolates whose cultivation was previously elusive perhaps because of the absence of their required specific plant polysaccharides. The human genome encodes enzymes that break down only starch, sucrose, and lactose (El Kaoutari et al., 2013). Carbohydrates make up 45-65% of typical human diets (Board., 2005). Therefore, the ability of intestinal microbes to break down polysaccharides is critical for human survival. Whether *P. copri* strain-type and presence is meaningful to human health, or simply an indication of diet, requires further investigation. Future studies that aim to assess human microbiome interactions with disease will require dietary intervention or, at minimum, collection of patient dietary habits or records. To avoid confounding effects of diet, future metagenomic analyses may better illuminate disease-associated genes by excluding genes known to interact with host diet. If *P. copri* does exacerbate or ameliorate disease in specific populations, diet may be readily manipulated to decrease or increase *P. copri* growth. Establishment of polysaccharide substrate preferences for genetically diverse human *P. copri* isolates allows for informed experimentation *in vitro* and *in vivo*, as well as clinical study design and interpretation.

## Acknowledgements

We thank Paul Zappile and Adriana Heguy at the NYU Genome Technology Center for sequencing support. We thank Rhina Medina and Lucy Alvarado for sample collection, Michael Fischbach for *Bacteroides* strains, Gretchen Diehl for critical reading of the manuscript, and Dawn Hershey for helpful discussions about dietary fiber. We thank Curtis Huttenhower for helpful discussions and for connecting us with NS and AT.

The NYU Genome Technology Center was partially supported by the Cancer Center Support Grant P30CA016087. HF-P was supported by an NYU-HCC CTSI grant (1TL1TR001447). JUS is funded through NIH/NIAMS R01AR074500. JUS is further supported by The Riley Family Foundation, The Beatriz Snyder Foundation, the Rheumatology Research Foundation and The Judith and Stewart Colton Center for Autoimmunity. LMC was supported by NIH Grant 1UL1RR029893. NS was supported by European Research Council (ERC-STG project MetaPG-716575). RB was supported by the Simons Foundation. DRL was supported by the Howard Hughes Medical Institute, the Colton Center for Autoimmunity, and the Rainin Foundation.

## Author contributions

HF-P, VR, and CM designed the study with input from DRL and RB. HF-P isolated *Prevotella* and designed and performed all experiments, with isolation strategy by LMC. JUS provided stool samples from NORA patients. CM analyzed 16S sequencing data and genomes. VR predicted PULs from the isolate genomes and helped design growth experiments. AT and NS identified *susC* clusters in metagenomic data. CG provided technical assistance for growth experiments and analysis of PULs. AW and JDW-G helped analyze genome and growth data. HF-P wrote the manuscript with contributions from VR, CM, and DRL, and input from all other authors.

## STAR Methods

### KEY RESOURCES TABLE

**STAR Methods:** **KEY RESOURCES TABLE**
| REAGENT or RESOURCE | SOURCE | IDENTIFIER |
| --- | --- | --- |
| Biological Samples |  |  |
| Fecal samples from new onset rheumatoid arthritis patients and healthy controls | This study | N/A |
| Critical Commercial Assays |  |  |
| PowerSoil DNA isolation kit | Qiagen | Cat No./ID: 12888-100 |
| TruSeq DNA PCR-free Library Prep Kit | Illumina, California, USA | 20015962 |
| Wheaton™ CryoELITE™ Cryogenic Storage Vials | Fisher Scientific | 02-912-737 |
| FALCON 14 ml Polypropylene Round-Bottom Tubes | Fisher Scientific | 352059 |
| Titertube® Micro Test Tubes | Bio-Rad | 2239391 |
| Deposited Data |  |  |
| *P. copri* isolate genomes | This study | NCBI-BioProject: **PRJNA559898** and PATRIC: https://www.patricbrc.org/ |
| 16S sequencing | This study | NCBI-BioProject: **PRJNA559898** |
| Experimental Models: Organisms/Strains |  |  |
| *Prevotella copri* DSM 18205 | DSMZ | DSM 18205 |
| *Bacteroides ovatus* ATCC8482 | Michael Fischbach |  |
| *Bacteroides thetaiotaomicron* VPI-5482 | Michael Fischbach | N/A |
| *P. copri* isolates | This study | N/A |
| Oligonucleotides |  |  |
| Primers for *Prevotella* and *P. copri* screen, see **Table S1** | This study | N/A |
| Software and Algorithms |  |  |
| DADA2 | Callahan et al., 2016 | https://benjjneb.github.io/dada2/ |
| Trimmomatic | Bolger et al., 2014 | http://www.usadellab.org/cms/?page=trimmomatic |
| SPAdes | Bankevich et al., 2012 | http://cab.spbu.ru/software/spades/ |
| PanPhlAn | Scholz et al., 2016 | https://bitbucket.org/CibioCM/panphlan/src/default/ |
| CheckM | Parks et al., 2015 | https://ecogenomics.github.io/CheckM/ |
| PATRIC | (Wattam et al., 2014) | https://www.patricbrc.org/ |
| MUSCLE | Edgar, 2004 | https://www.drive5.com/muscle/ |
| RAxML | Stamatakis, 2014 | https://cme.h-its.org/exelixis/web/software/raxml/index.html |
| Silva SINA aligner | (Pruesse et al., 2012) | https://www.arb-silva.de/aligner/ |
| Other |  |  |
| Anaerobic Tissue Transport Medium Surgery Pack – ATTMSP | Anaerobe Systems | AS-915 |
| Brucella Blood Agar - BRU | Anaerobe Systems | AS-141 |
| Laked Brucella Blood Agar w/ Kanamycin and Vancomycin – LKV | Anaerobe Systems | AS-112 |
| Peptone Yeast Broth with Glucose - PYG | Anaerobe Systems | AS-822 |
| Yeast Casitone Fatty Acids Broth with Carbohydrates – YCFAC Broth | Anaerobe Systems | AS-680 |
| Polysaccharides, see Table S4 | see Table S4 | see Table S4 |

### LEAD CONTACT AND MATERIALS AVAILABILITY

Further information and requests for resources and reagents should be directed to and will be fulfilled by the Lead Contact, Dan R. Littman.

### EXPERIMENTAL MODEL AND SUBJECT DETAILS

Subjects enrolled in this study included new onset rheumatoid arthritis patients and healthy controls, as described in STAR methods. Ethical approval was granted by the Institutional Review Board of New York University School of Medicine.

### METHOD DETAILS

#### Sample collection

Stool was collected into anaerobic collection tubes (Anaerobe Systems) from healthy controls and new onset rheumatoid arthritis patients, deidentified, and frozen at −80°C. New onset rheumatoid arthritis patients were untreated, presenting with symptoms, seropositive for anti-citrullinated protein antibodies, and not on antibiotics in the previous six months.

#### qPCR Screen for *Prevotella*

Genomic fecal DNA was extracted from fecal samples using the MoBio PowerSoil DNA extraction kit. Fecal DNA was screened for the presence of *P. copri* using qPCR primers designed for six conserved regions of the *P. copri* genome (Scher et al., 2013), as well as two conserved regions of the *Prevotella* 16S gene (Table S1). Universal primers specific for the 16S rRNA gene were used to quantify total bacterial load in each sample. qPCR was performed with the SYBR green master mix on the Roche Lightcycler 480, with the following cycle conditions: 90°C for 5 minutes, then 40 cycles of 95°C for 10 seconds, 60°C for 30 seconds, and 72°C for 30 seconds. Genomic DNA from *P. copri* DSM 18205 was used to generate a standard curve. The standard curve was used to determine the absolute quantity of *Prevotella* and 16S in a sample. Fold change was calculated by dividing absolute quantity of *Prevotella* by absolute quantity of 16S. Feces with a fold change > 0.1 for any *Prevotella*-specific primer set were considered positive and used to isolate *P. copri*, with the exception of sample AU, which had a lower fold change.

#### Isolation of *Prevotella*

*P. copri* was isolated from *Prevotella*-positive feces by anaerobic quadrant streaking of PBS-diluted and undiluted stool on BRU and LKV plates (Anaerobe Systems), then incubated anaerobically at 37°C for 24-48 h. Isolate colonies were picked, enumerated, streaked on a fresh plate, and the inoculation loop dipped into PCR grade water. The loop-dipped PCR water was used with the qPCR primer sets (above) to screen for *P. copri*. *Prevotella*-positive-screened isolates were confirmed by Sanger sequencing of the V3-V4 hypervariable region of the 16S rRNA gene, and finding >97% identity to known *Prevotella* species. Glycerol stocks of *Prevotella* isolates were frozen at −80°C.

#### 16S rRNA gene sequencing

Genomic fecal DNA was extracted from fecal samples using the MoBio PowerSoil DNA extraction kit. TheV3-V4 region of the 16S rRNA gene was amplified, purified, and sequenced on Illumina MiSeq (Caporaso et al., 2012). Sequences generated from amplicon sequencing were trimmed (maxN=0, truncQ=2) and denoised using DADA2 (Callahan et al., 2016). Sequences were annotated using the Silva nr v128 database.

#### Whole genome sequencing

Isolates were grown on BRU plates for 24-48 hours, until a healthy lawn was present. The lawn was suspended in PYG media (Anaerobe Systems), pelleted, and DNA was isolated with the MoBio PowerSoil DNA isolation kit. Libraries were prepared for sequencing with the TruSeq PCR-free library preparation kit and sheared to 500bp fragment length. Prepared libraries were sequenced on Illumina HiSeq2500 2×100 bp. Paired-end sequences were trimmed using Trimmomatic (Bolger et al., 2014), then whole genomes were assembled using SPAdes (Bankevich et al., 2012). Trimmed reads were also mapped to the panphlan_pcopri_16 database (https://bitbucket.org/CibioCM/panphlan/wiki/Pangenome%20databases) using PanPhlAn (default parameters, Figure 2A) (Scholz et al., 2016). Quality and completeness of the assemblies were evaluated using CheckM (Parks et al., 2015). Genomes were annotated in PATRIC and are available at https://www.patricbrc.org/ and under BioProject **PRJNA559898**. Presence and absence of predicted proteins can be visualized at https://flatironinstitute.github.io/genome_cluster_array_visualization/.

#### Phylogenetic Trees

Representative genomes for the *Prevotella* and *Bacteroides* genera were downloaded from NCBI. For each genome, a set of 30 ribosomal protein genes (LSU ribosomal proteins L1, L2, L3, L4, L5, L6, L10, L13, L14, L15, L18, L22, L23, L24, L29, and SSU ribosomal proteins S2, S3, S4, S5, S7, S8, S9, S10, S11, S12, S13, S14, S15, S17, S19) were identified and aligned using MUSCLE (Edgar, 2004). A maximum-likelihood tree was calculated using RAxML (-m GTRCAT; (Stamatakis, 2014)) from a concatenated alignment of the 30 ribosomal protein genes. A 16S rRNA gene tree was also estimated from the alignment of full length 16S rRNA gene sequences using Silva’s SINA aligner (https://www.arb-silva.de/aligner/) and RAxML (-mGTRCAT).

#### *SusC* analysis and polysaccharide utilization prediction

All annotated *susC* genes in the isolate genomes were clustered at >90% identity. This analysis revealed 168 *susC* gene clusters throughout all the genomes (Figure S2). For each *susC* gene cluster, ten flanking genes on either side were pulled from the genome and annotated (Table S3). Flanking genes were examined and classified according to predicted substrate. *SusC* clusters lacking a companion *susD* gene, as well as those comprised of genes unrelated to polysaccharide catabolism, were excluded from the pPUL repertoire analysis.

#### *In vitro* polysaccharide growth assays

*B. thetaiotaomicron*, *B. ovatus* (both from Michael Fischbach), *Prevotella copri* DSM 18205 (DSMZ), and *P. copri* isolates were grown anaerobically on BRU plates (Anaerobe Systems) for 48h at 37 degrees C. Strains were passaged to YCFAC liquid media (Anaerobe Systems). 48h later, cultures were passaged to pre-reduced YCFA +/- 0.5% (w/v) individual polysaccharides (Table S4) at a dilution of 1:100 in a total volume of 1ml (Browne et al., 2016). Cultures were incubated anaerobically in sterile polypropylene tubes at 37 degrees C. Importantly, P. copri grew in Wheaton™ CryoELITE™ Cryogenic Storage Vials, FALCON 14 ml Polypropylene Round-Bottom Tubes, and Titertube® Micro Test Tubes (see **Key Resources Table**), but not in Axygen™ Storage Microplates from Fisher Scientific (Catalog no. 14-222-224). Every 24h, 100 μl of each culture was transferred to a flat-bottom 96-well plate, and OD600 was measured with the Envision plate reader. OD600 for each culture condition was subtracted from the average blank OD600 for the corresponding polysaccharide. Cultures were grown in duplicate or triplicate. For each condition, the maximum average OD600 from the 24h, 48h, or 72h time point was determined and duplicate or triplicate values were used in Figure 4A, 4B, and Figure S5.

#### PUL prediction in *P. copri* MAGs

*SusC* and *susD* homologs in *P. copri* MAGs (Tett et al., 2019) were identified by blastp (Identity>75%, Bitscore > 300) using *susC* and *susD* predicted proteins from the *P. copri* isolate genomes. Only MAG contigs containing both *susC* and *susD* homologs within 10 gene products were considered further. For each *susC* centroid, the PUL operon was extended to include the 10 gene products upstream and downstream of the first and last occurrence of a *susC* or *SusD* homolog. Each PUL was checked for CAZy annotations (Lombard et al., 2014) predicted using HMMSEARCH (version 3.1b2) (Eddy, 2011) against dbCAN HMMs v6 (Yin et al., 2012) with an E-value < 1e-18 and coverage > 0.35. PULs were predicted by searching for the presence of CAZymes known to act on known polysaccharides. Predictions are presented as the proportion of *P. copri* MAGs containing at least one pPUL out of all *P. copri* MAGs within each *P. copri* complex clade.

### QUANTIFICATION AND STATISTICAL ANALYSIS

#### Prediction analysis

For each isolate used in the *in vitro* polysaccharide growth assay, a one-way ANOVA was performed on the maximum OD for each polysaccharide. A post hoc Dunnet’s test was performed with the H2O condition (no carbon source) as the negative control. Isolate-polysaccharide conditions with OD values that were statistically significant compared to the H2O control were considered positive for growth. Isolate-polysaccharide conditions that were not significant were considered negative for growth.

#### Statistical analysis of prediction

Isolate-polysaccharide conditions were separated according to whether growth was predicted: if a pPUL was present in the genome of an isolate, the isolate-polysaccharide condition was considered “growth predicted.” If a pPUL was absent in the genome of an isolate, the isolate-polysaccharide condition was considered “growth not predicted.” Isolate-polysaccharide conditions that were statistically significant as determined by a one-way ANOVA with a post hoc Dunnet’s test using the H2O condition (no carbon source) as the negative control were given a score of 1 and isolate-polysaccharide conditions that were not significant were given a score of 0. Welch’s t test was performed on the data.

## DATA AVAILABILITY

All 16S sequencing data and whole genome sequencing data are available under BioProject PRJNA559898.

## Supplemental Information

**Figure S1.**
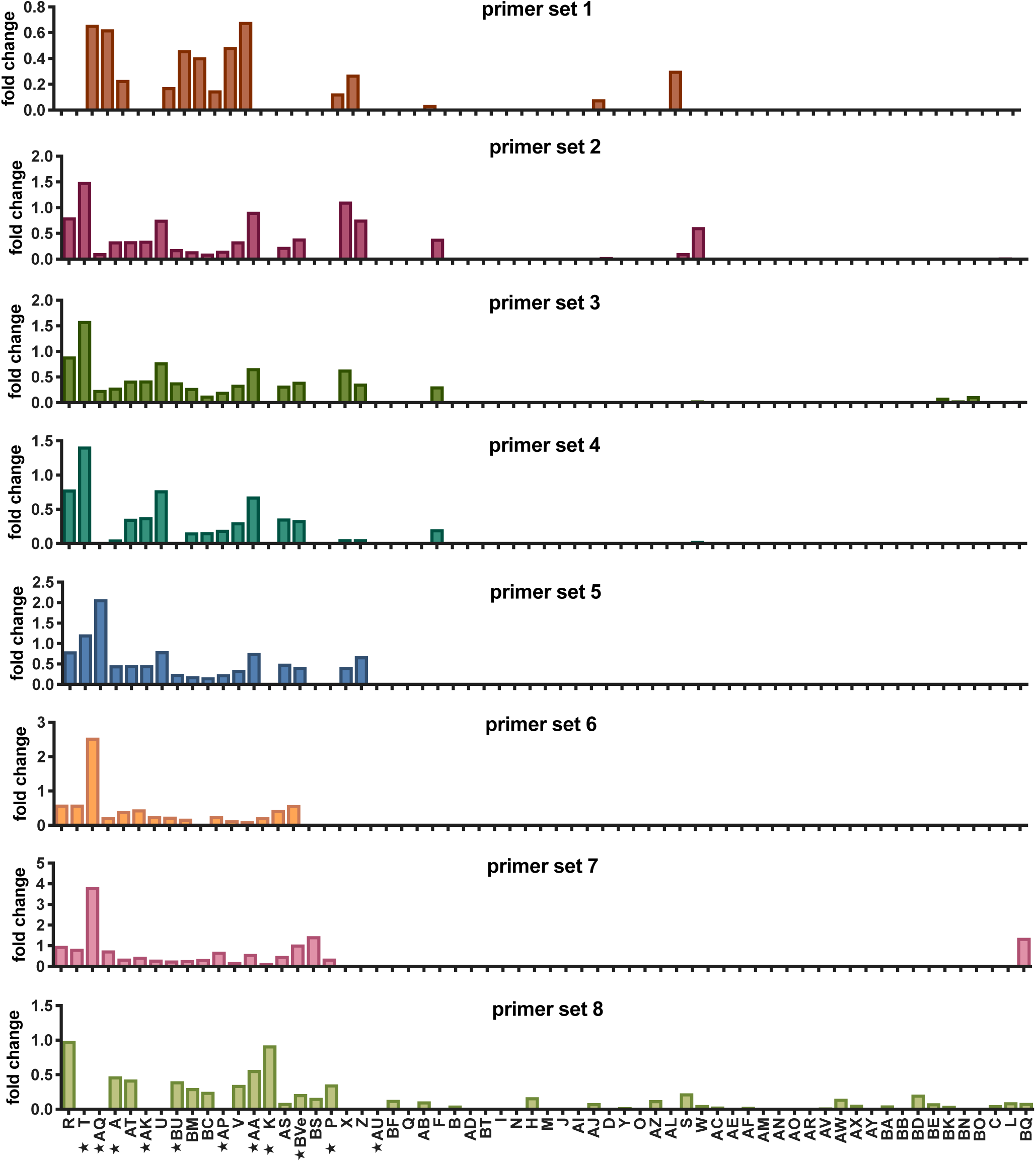
PCR-based screen for *P. copri*-containing fecal microbiota; related to Figure 1. Eight sets of primers were designed to amplify conserved regions of the *P. copri* genome. Fecal DNA from each donor was screened by qPCR using each primer set (Table S1). Stars indicate individuals from whom *P. copri* was isolated.

**Figure S2.**
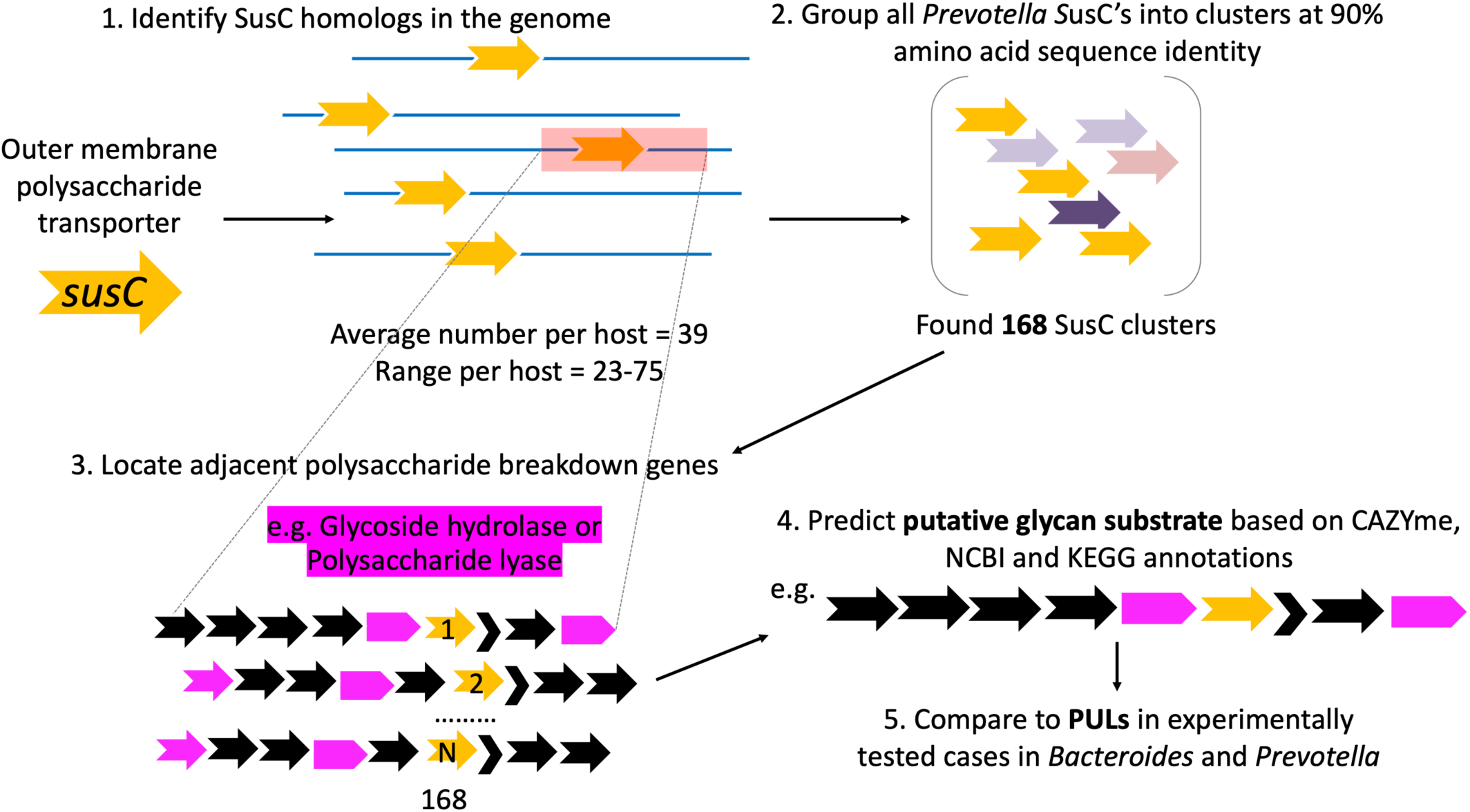
Identification of polysaccharide utilization loci in *P. copri* isolates; related to Figure 3. *SusC* homologs were identified in the genomes and clustered by 90% sequence identity. Adjacent polysaccharide genes were identified and used to annotate putative PULs from each *susC* cluster.

**Figure S3.**
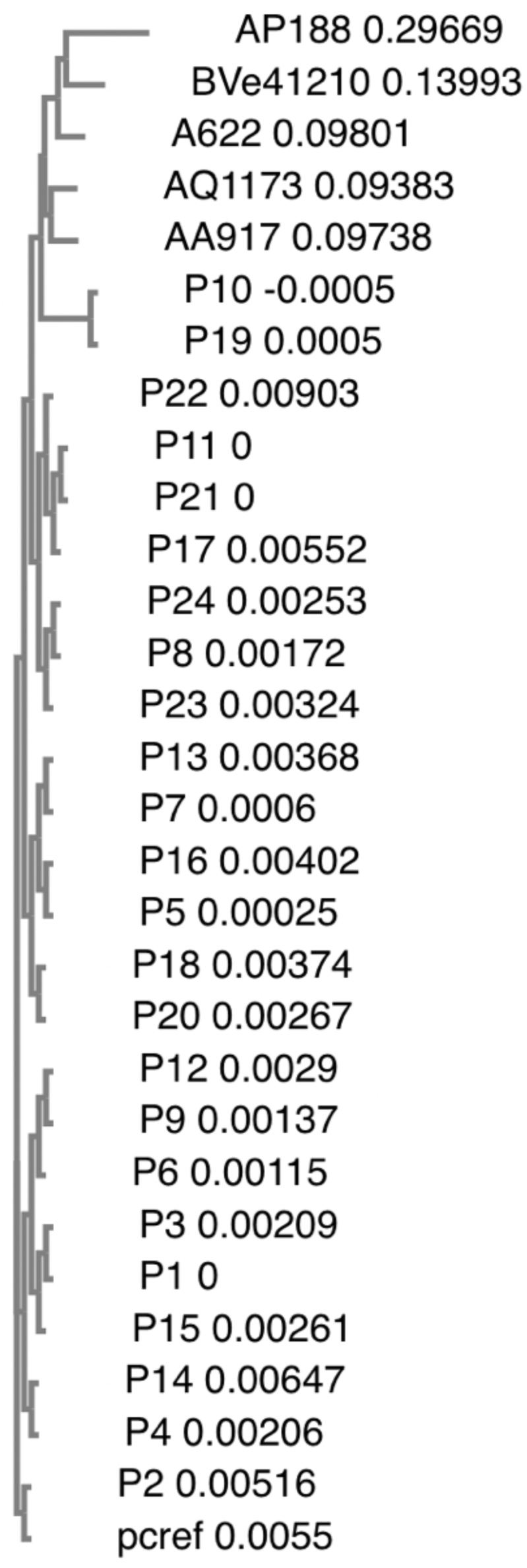
*Prevotella* 16S V1-V3 sequences; related to Figure 3. Representative isolates were selected and the full length 16S sequence was determined by Sanger sequencing. 16S hypervariable V1-V3 sequences were aligned with *Prevotella* 16S V1-V3 sequences described in De Filippis, et al. (2016) using Clustal Omega. A simple phylogeny is shown.

**Figure S4.**
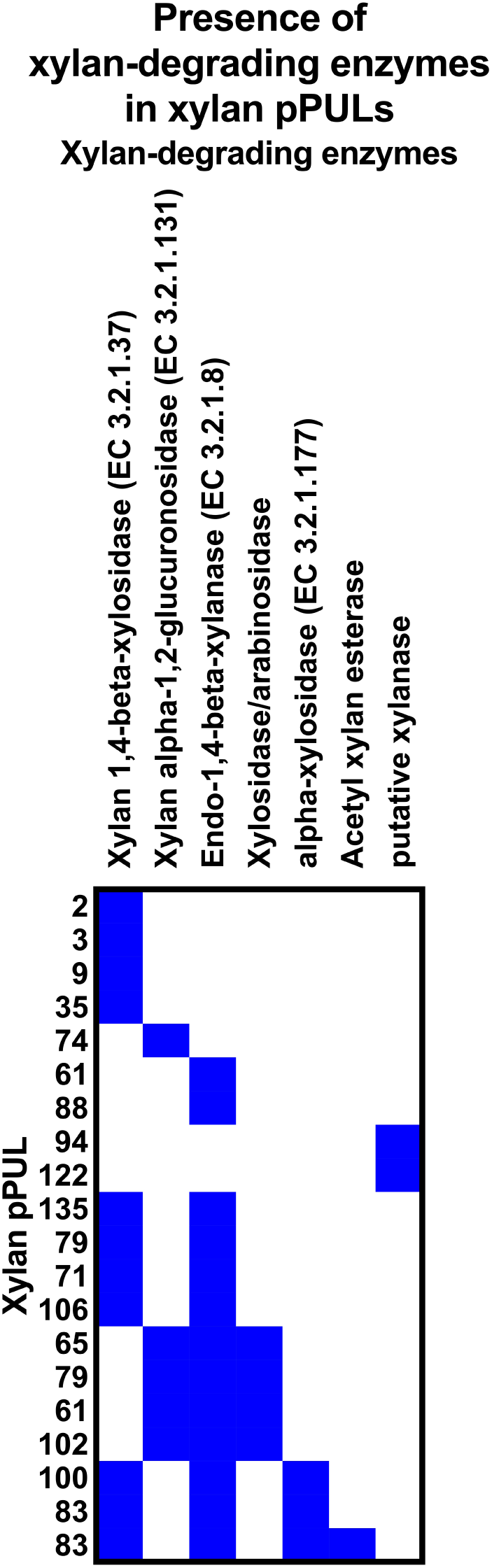
Xylan enzyme distribution in xylan pPULs; related to Figure 3. Xylan pPULs and xylan-degrading enzyme annotations are shown. Presence of each xylan-degrading enzyme in each pPUL is indicated by a blue box.

**Figure S5.**
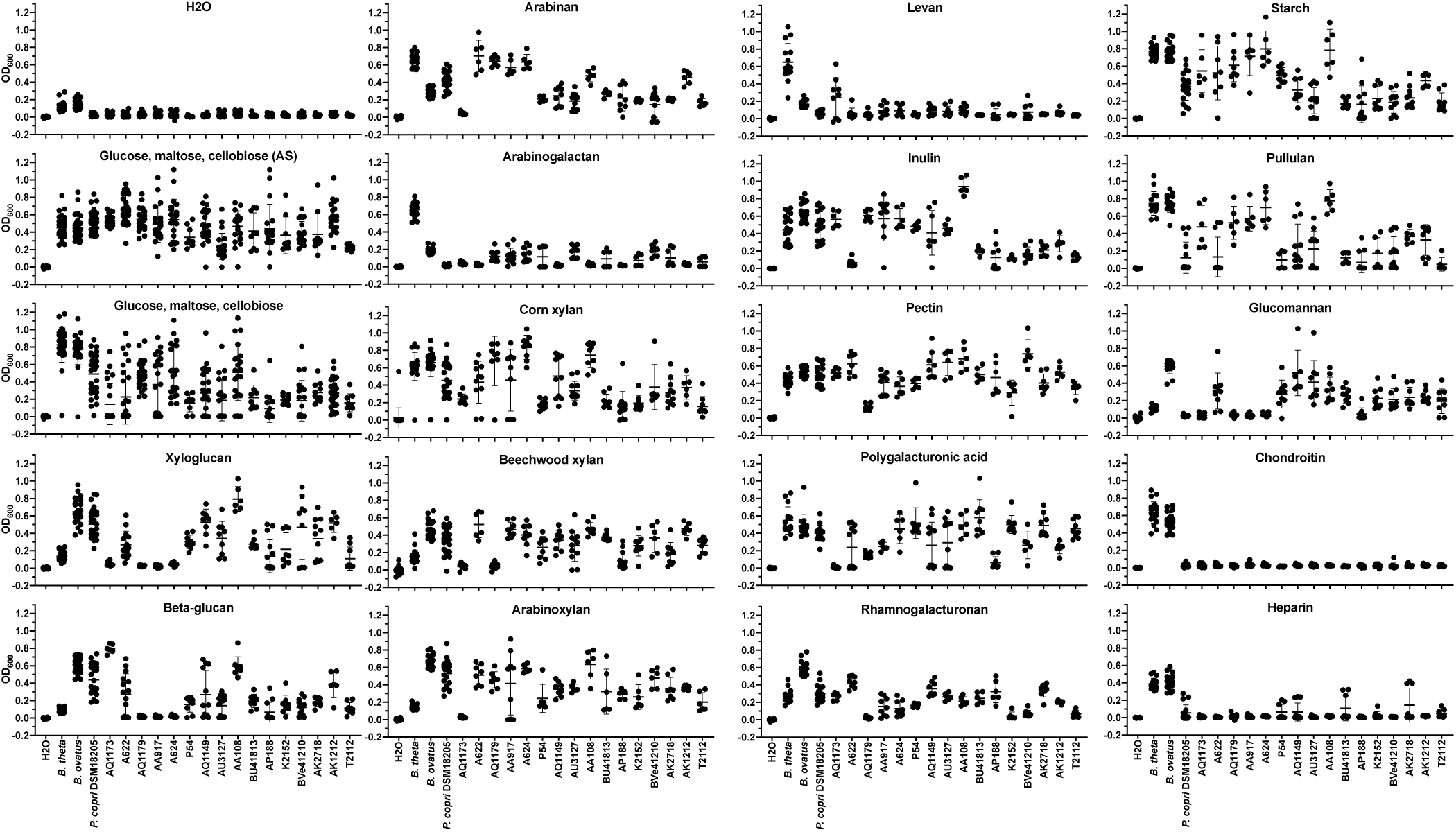
Growth of *P. copri* isolates on polysaccharides; related to Figure 4. OD600 was measured for indicated isolates and polysaccharides after ∼24h, ∼48h, ∼72h anaerobic incubation at 37 degrees C. Data represent 2-3 experiments. Each condition was tested in duplicate or triplicate. Data show the highest OD from 24, 48, or 72h for each condition for each experiment.

## Supplemental Tables

**Table S1.**
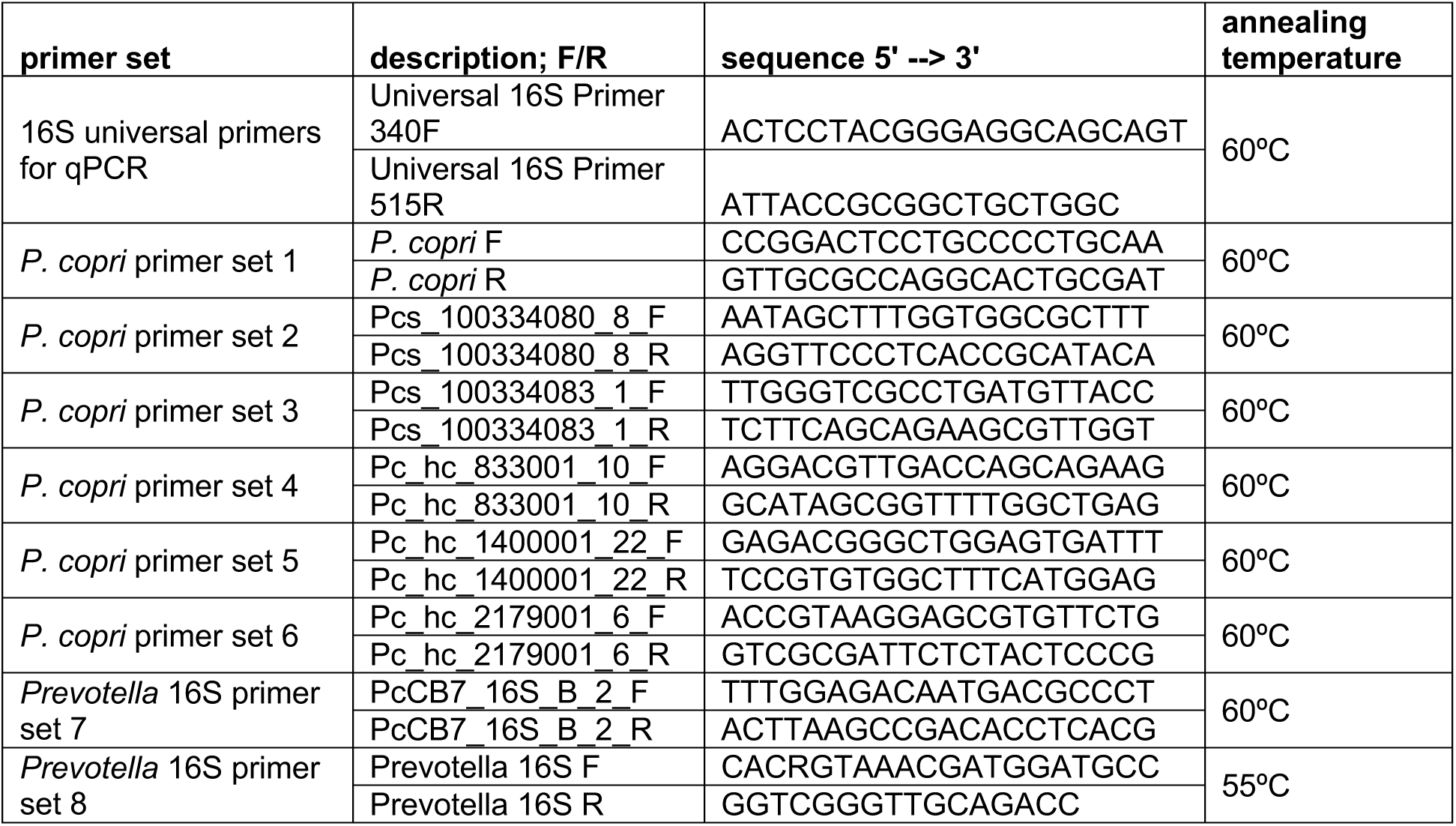
*P. copri* primers table, related to Figure 1 and Figure S1.

**Table S2. CheckM summary, related to Figure 2**

**Table S3. *SusC* gene clusters, related to Figure 3**

**Table S4.** Polysaccharides used, related to Figure 4.

| **Polysaccharide** | **Company** | **Catalog number** |
| --- | --- | --- |
| Arabinan (Sugar Beet) | Megazyme | P-ARAB |
| Arabinogalactan (Larch Wood) | Megazyme | P-ARGAL |
| Arabinoxylan (Wheat Flour; Low Viscosity ~ 10 cSt) | Megazyme | P-WAXYL |
| Beta-glucan (Barley; High Viscosity) | Megazyme | P-BGBH |
| Chondroitin sulfate sodium salt (from shark cartilage) | Sigma | C4384-5G |
| Glucomannan (Konjac; Low Viscosity) | Megazyme | P-GLCML |
| Heparin sodium salt (from porcine intestinal mucosa) | Sigma | H3393-100KU |
| Inulin (chicory) | Megazyme | P-INUL |
| Levan (from *Erwinia herbicola*) | Sigma | L8647-1G |
| Pectin (from apple) | Sigma | 93854-100G |
| Polygalacturonic Acid (PGA) | Megazyme | P-PGACT |
| Pullulan (from *Aureobasidium pullulans*) | Sigma | P4516-1G |
| Rhamnogalacturonan (Soy Bean) | Megazyme | P-RHAGN |
| Starch | VWR | 1.01252.0100 |
| Xylan (Beechwood; purified) | Megazyme | P-XYLNBE-10G |
| Xylan (from Corn Core, TCI America™) | Fisher | X007825G |
| Xyloglucan (Tamarind) | Megazyme | P-XYGLN |

## Supplemental Files

File S1. *P. copri* 16S SNV sequences, related to Figure 1

File S2. *P. copri* 16S V1-V3 sequences aligned to *P. copri* 16S sequences from De Filippis et al., 2016, related to Figure 3

